# Modular chronic cranial platform for longitudinal functional ultrasound, optical, and ECoG recording in a gyrencephalic model species

**DOI:** 10.64898/2026.08.24.746649

**Authors:** Klaudia Csikós, Ábel Petik, Domonkos Horváth, Attila Balázs Dobos, Ágoston Horváth, Dries Kil, Alan Urban, Botond Roska, Daniel Hillier

## Abstract

Understanding cortical function in large, gyrencephalic brains requires following the same tissue over time and interrogating it through complementary methods — yet in practice each recording modality is run on its own preparation, across separate animals and sessions that cannot be quantitatively related, while the recorded cortex can rarely be revisited. Functional ultrasound (fUS), widefield optical imaging, and electrocorticography (ECoG) are individually mature and strongly complementary, but no single preparation has combined all three within the same chronically accessible tissue in a gyrencephalic brain. Here we present a modular chronic cranial platform, validated in three cats with implants remaining functional for up to 3.3 years that unites these modalities in one customized chamber built around fUS as an anchor modality. The platform also supports fUS imaging in awake, head-unrestrained animals, with activation maps remaining spatially consistent across imaging days. By providing stable, quantitatively reproducible access to the same cortical region over time, this platform enables longitudinal, multimodal characterization of cortical function within individual subjects.

## Introduction

Brain function is organized across spatial and temporal scales that no single recording method resolves at once — from individual neurons on the millisecond timescale, through local circuits spanning hundreds of micrometers to millimeters, to distributed cortical networks extending across centimeters and evolving over seconds (Munn et al., 2024; Higley & Cardin, 2022). Effort has concentrated on individual modalities and their resolution limits, yet in large, gyrencephalic brains the more severe limitation is not the sensitivity of any one method, but the fragmentation of the experimental program built around it. A typical program is split across separate animals — one cohort imaged, another recorded — across sessions that cannot be related to one another quantitatively. This fragmentation carries three costs. First, between-animal and between-session variance becomes confounded with the effect of interest, so pooled results are fragile and difficult to reproduce. Second, any process that unfolds over time — plasticity, learning, adaptation, recovery — is invisible to a design that samples each animal only once. Third, the tissue itself remains out of reach: a preparation built for a single acute recording cannot later be returned for targeted perturbation.

What is missing therefore is not another modality but a platform that removes these constraints simultaneously (Ramezani et al., 2024). Four properties define it: (i) chronic, repeated access to the same cortical location, the prerequisite for following any process over time thereby accumulating within-animal statistical power; (ii) reproducible quantitative data across sessions, so that a difference between two timepoints is a measurement rather than an artifact of spatial misalignment; (iii) interchangeable modalities on one preparation, so that complementary readouts come from the same tissue without a new surgery for each; and (iv) physical access for intervention, so that functional recording and perturbation can be run on one animal across the lifetime of one implant. Chronic, reproducible access is the load-bearing property: without it, multimodality collapses into a set of unrelatable single sessions, and longitudinal questions cannot be posed at all.

Functional ultrasound (fUS) is well suited to anchor such a platform. Operating natively at the mesoscale, fUS measures stimulus-evoked changes in cerebral blood volume across centimeters of tissue and to centimeter depths, with spatial resolution approaching 100 μm and temporal resolution of ∼100 ms to a few hundred milliseconds depending on imaging depth (Macé et al., 2011; Tanter & Fink, 2014; Deffieux et al., 2018). Crucially for persistent access, fUS images vasculature — a dense, animal-specific anatomical scaffold that provides a natural substrate for locating the same cortical volume across sessions. Chronic fUS through a cranial window is established in rodents (Urban et al., 2014, 2015; Macé et al., 2018) and has been extended to gyrencephalic brains — the awake ferret (Bimbard et al., 2018) and the behaving macaque (Dizeux et al., 2019; Blaize et al., 2020) — although chronic, longitudinal preparations remain comparatively rare at this scale. fUS does require a craniotomy for high-signal-to-noise access, since the skull strongly attenuates and aberrates ultrasound; but this is a one-time cost paid at the level of the skull, not the brain: once the window is in place, the same tissue can be revisited across months without the cumulative cortical damage that accompanies repeated electrode insertion or optical window reimplantation.

The other two modalities extend what fUS alone provides. Widefield intrinsic signal optical imaging offers a well-established framework for mapping functional cortical organization on the surface and for validating mesoscale activity patterns measured with fUS (Grinvald et al., 1986; Bonhoeffer & Grinvald, 1991), and the same optical port opens a route to multi-photon imaging of the cellular substrate of mesoscale dynamics (Schummers et al., 2008). Conversely, fUS extends optical imaging by resolving the full cortical thickness and, depending on the imaging depth, subcortical structures beyond the largely surface-confined reach of optics. ECoG complements both imaging modalities by providing a concurrent measure of cortical electrical activity, enabling hemodynamic and metabolic signals to be related directly to their underlying electrophysiological dynamics and changes in brain state.

Each pairwise combination has precedent — fUS with electrophysiology (Sieu et al., 2015; Bergel et al., 2018, 2020; Claron et al., 2023; Panskus et al., 2026), fUS with two-photon microscopy in the same animal (Boido et al., 2019), and ECoG with optical imaging through a transparent “smart dura” at non-human-primate scale (Yazdan-Shahmorad et al., 2016; Griggs et al., 2021) — but rarely in a large, gyrencephalic brain, and never as a single chronic multimodal platform integrating all three modalities over the same tissue. The only demonstration of the full triad, the COMBO window (Edelman et al., 2024), was built for the lissencephalic mouse.

Longitudinal access is not a convenience but the precondition for whole classes of questions: following plasticity, separating genuine biological change from session-to-session variation in anesthesia or arousal, and building within-animal statistical power without inflating animal numbers. Yet “chronic” is more often asserted than measured — a preparation is called chronic because it survives, not because the same tissue has been shown to be reproducibly identified and sampled across the intended timescale. For a platform whose central promise is long-term accessibility, that promise must be demonstrated as a quantity: how accurately, and how stably over months, a previously recorded cortical volume can be found again. In terms of functional imaging, reproducibility across sessions further depends on an anesthesia protocol whose vascular and neurovascular-coupling effects are compatible with a cerebral blood volume (CBV)-based readout (Nunez-Elizalde et al., 2022); because established fMRI anesthesia protocols cannot be assumed to transfer to fUS, we developed and separately validated an optimized isoflurane–ketamine–medetomidine protocol for this purpose (Horváth et al., companion paper), used throughout the anesthetized recordings here.

Here we present a modular chronic cranial chamber and head-fixation system, validated in the cat, designed around multi-month, repeatable access to the same cortical tissue in the same animal. A single chamber body accepts swappable fUS and optical inserts combined with ECoG. An MRI-based, individual-animal fitting pipeline conforms the geometry to each skull before surgery and predicts accessible areas of the cortex, which we confirm in vivo. We then established a vasculature-based registration pipeline that aligns a previously recorded cortical volume across experimental days with submillimeter precision. Anchored on this fUS-based registration, we demonstrate multimodal reach (concurrent fUS–ECoG; fUS-versus-optical imaging) and show that awake fUS acquisition without head fixation is reproducible across weeks. The cat provides a gyrencephalic cortex with a mature, well-characterized visual system and decades of established protocols, at higher throughput and lower cost than primate work, so that the validated design transfers to non-human primates as a principled dimensional adaptation rather than a redesign.

## Methods

### Animals and ethics

Three adult domestic cats (two females and one male, 2–4 years old) were used in this study. All animals were bred in the animal facility of the HUN-REN Research Centre for Natural Sciences, Budapest, Hungary. All surgical and experimental procedures were approved by the Animal Care Committee of the HUN-REN Research Centre for Natural Sciences and by the National Food Chain Safety Office of Hungary and were conducted in accordance with applicable regulations governing the care and use of experimental animals.

### Chamber design and fabrication

#### Individual-animal design pipeline

Each chamber is planned from a volumetric T1-weighted MRI scan of the target animal’s head (3T Siemens MAGNETOM Prisma), acquired with a T1 VIBE sequence tailored to resolve the skull surface at sub-millimeter precision alongside soft tissue, avoiding the need for a separate CT acquisition. Skull and brain are segmented automatically from the T1 volume (Fig. S1A), and the chamber footprint is defined by virtual skull cutting against the reference geometry (Fig. S1B). Predicted cortical access is then established by registering a 3D cat cortical atlas to the individual MRI image in 3D Slicer, allowing visual, auditory, and somatosensory areas within the planned craniotomy to be identified before surgery (Fig. S1C) (Stolzberg et al., 2017; Kikinis et al., 2014).

#### Modular architecture

Three interoperating components (Fig. 1A): (i) a chronic chamber whose skirt is contoured to the individual skull and whose upper surface presents a standardized docking geometry; (ii) a load-bearing metal headplate mounting onto that geometry, accepting extensions and connecting to head fixation (Fig. S2A); and (iii) swappable insets — a polymethylpentene (PMP) foil inset for fUS and a glass inset for widefield imaging — docking onto the same interface, so modalities are exchanged within one craniotomy window without further surgery (Fig. S2B–D).

**Figure 1.**
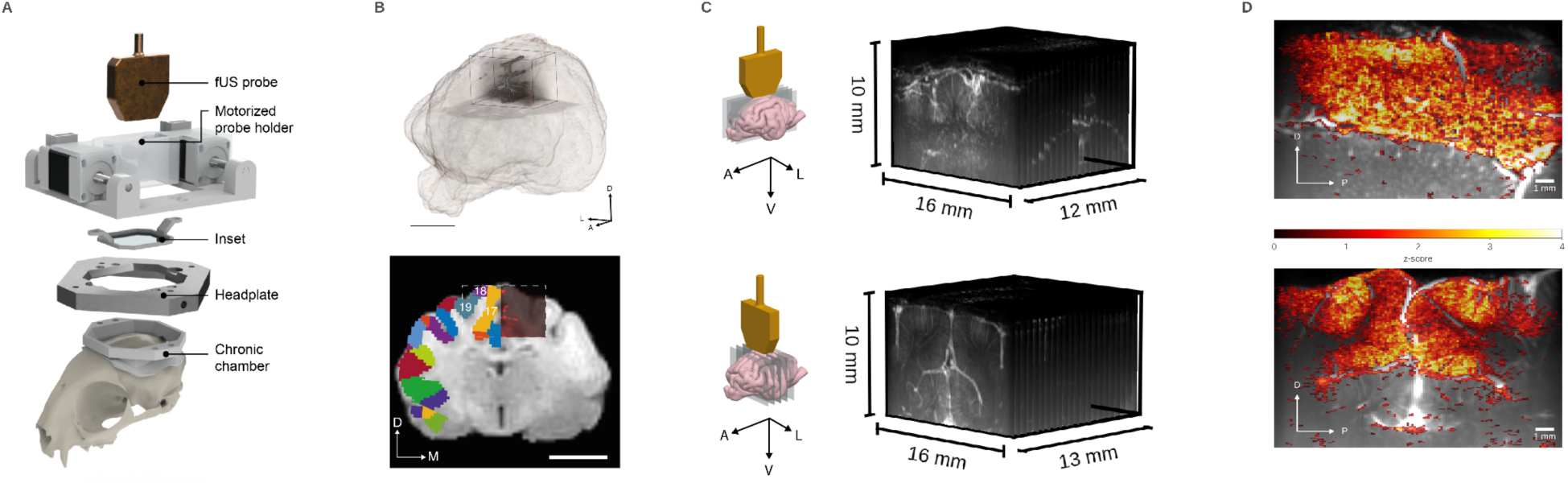
Modular implants for large-scale chronic fUS. (A) Exploded view of the modular implant system. From bottom to top: chronic chamber body on a cat skull model, load-bearing metal headplate, stabilizing inset with PMP foil, the motorized probe holder and the 12 MHz ultrasound transducer. (B) Co-registration of fUS and structural MRI volumes acquired from the same animal. (Top) 3D rendering showing the fUS imaging volume (dark cuboid) registered within the whole-brain MRI (semi-transparent), aligned manually using anatomical landmarks. (Bottom) Coronal section through the co-registered fUS and MRI volumes; the white dashed outline delineates the full extent of the fUS acquisition volume, which is shown “cut open” on the right half to reveal the underlying vasculature, while the left half displays the corresponding MRI slice overlaid with functional area labels from the co-registered cat cortical atlas (Stolzberg et al., 2017). The fUS volume encompasses visual areas 17, 18, and 19, confirming coverage of visual cortex. (C) In-vivo 3D power-Doppler scans acquired through the motorized fUS module along sagittal (top) and coronal (bottom) sweep axes (device-orientation illustrations, left), yielding vascular volumes of 16 × 10 × 12 mm and 16 × 10 × 13 mm respectively, acquired with 200 μm stepsize. (D) Stimulus-evoked fUS activation: pixel-wise z-score activation maps overlaid on structural power-Doppler images, showing spatially localized, visual stimulus-locked responses in visual cortex. Scale bars (B) 1 cm, (D) 1 mm. A, anterior; D, dorsal; L, lateral; M, medial; P, posterior; V, ventral

#### Motorized fUS module

The linear transducer is translated along two orthogonal axes by miniature linear stepper actuators (10 μm/step, Nanotec LSA201S06-A-UECB-102) under open-source microcontroller control, extending planar fUS into volumetric acquisition along coronal and sagittal axes within the chamber footprint (Fig. 1A,C; Fig. S2A,B). The module adds <5% of body weight.

#### Materials and fabrication

The customized chamber bodies and interchangeable insets were fabricated by stereolithographic 3D printing using a Form 3B+ printer (Formlabs) and Biocompatible BioMed Black Resin (Formlabs). The motorized headplate was likewise 3D-printed using Rigid 4K Resin (Formlabs). The load-bearing metal headplate was CNC-machined from EN AW-6061 aluminum alloy. Stainless steel threaded inserts were heat-set into the 3D-printed chamber bodies to provide threaded mounting interfaces. Interchangeable tissue-stabilizing inner elements at fixed depth increments provide adjustable mechanical contact with the cortical surface, allowing physiological pulsation to be stabilized while minimizing tissue compression. The artificial dura consists of a thin, optically clear PDMS film, positioned at durotomy to provide a stable interface over the exposed cortical surface and limit dural regrowth.

### Surgical implantation and chronic maintenance

Implantation proceeds in three staged surgeries — outer chamber implantation, craniotomy, and durotomy with definitive artificial-dura placement — each separated by ≥2 weeks of recovery, confining every surgery to one task. All three surgeries use one anesthesia protocol (alfaxalone/ketamine/fentanyl induction, neuromuscular blockade with fully controlled ventilation, prophylactic anti-edema management), distinct from the shorter recording-session protocol below (Horváth et al., companion paper). During the initial surgery, the outer chamber was stereotaxically aligned, anchored using a dental etch-and-prime bonding system, and reinforced with bone cement and stainless steel screws distributed along the anteroposterior axis to ensure long-term stability and osseointegration. Hydroxyapatite paste was applied to all tissue-contacting cement surfaces to encourage soft-tissue integration. In the second surgery, a 30 x 30 mm craniotomy was performed using progressive, saline-irrigated drilling to prevent thermal injury; the underlying dura was preserved intact or minimally fenestrated before temporary sealing with silicone and a protective cap. In the final surgery, the durotomy was completed and the definitive artificial dura was positioned over the cortex. Post-operative care follows a standard analgesia/antibiotic course with defined emergence and monitoring criteria. Drug doses, infusion-rate preparation, intraoperative edema management, and post-operative schedules are in Supplementary Protocol S1.

At each imaging session (typically twice weekly per animal) the chamber was cleaned with sterile saline and inspected for leakage, mechanical integrity, and stability; revision was considered on team assessment of loosening or structural damage.

To maintain the integrity and cleanliness of the chamber–skin interface, routine wound care was performed three times per week. Each maintenance session consisted of inspection, cleaning and disinfection of the wound site, followed by application of a fresh protective dressing.

### Acquisition

#### Experimental conditions

Anesthetized sessions follow the isoflurane–ketamine–medetomidine protocol validated int the cat, with individualized, pharmacokinetically-guided medetomidine dosing (full protocol, comparison against alternative regimens, and PK/PD modeling: Horváth et al., companion paper). In awake sessions the animal was seated before the stimulation screen with the body gently restrained and the head unrestrained; no gaze or head fixation was used beyond attachment of the transducer module to the chamber.

#### Functional ultrasound

Power-Doppler data were acquired with a 128-element, 12 MHz linear array (Urban et al., 2015) on a Vantage 256 scanner (Verasonics) at 1 kHz pulse-repetition frequency, giving Doppler images at 5 Hz after plane-wave compounding and SVD clutter filtering (Macé et al., 2011; Tanter & Fink, 2014). The transducer was positioned over visual cortex coronally or sagittally: 16 × 10 mm (width × depth) field of view, 125 × 60 μm in-plane voxels, ∼400 μm slice thickness. Fixation screws of the motorized holder secure the transducer above the cortex, and the module steps the probe through predefined planes to reconstruct volumes across sessions (Fig. 1C). Post-implantation, a 3D power-Doppler scan of the cranial window was co-registered manually by anatomical landmarks in 3D Slicer to the animal’s own pre-implantation MRI, and thereby to CATLAS space, assigning vascular landmarks to atlas-defined regions (Fig. 1B).

#### Widefield optical imaging

A custom tandem macroscope pairs a 50 mm f/0.95 imaging lens (DO-5095, Navitar) with a 50 mm f/1.4 objective (Nikkor AI-S, Nikon) via a 60 mm filter cube (Thorlabs); equal focal lengths give unity magnification, a field of view equal to the sensor (13.3 × 13.3 mm), 40 mm working distance, and effective NA 0.36. Images of cortical reflectance were obtained through a 525/50 nm bandpass filter (#86-963, Edmund Optics), close to an isosbestic point of oxy- and deoxyhemoglobin, measuring total hemoglobin concentration changes. Images were recorded by an sCMOS camera (Prime BSI Express, Teledyne Photometrics; 2048 × 2048, 6.5 μm pixels). Acquisition used 2×2 spatial binning (13 μm effective pixel) at 22 fps with 44 ms exposure, and two-frame temporal binning for an effective 11 fps.

#### Electrocorticography

An acute ECoG surface array was placed on the cortex under the inset window during fUS recordings, over the marginal gyrus (area 17) beneath the transducer (Fig. 3A; Fig. S2C). The device is a flexible polyimide microECoG with platinum sites and interconnects, 32 sites (150 μm) in a 4 × 8 grid at 500 μm spacing, covering 2 × 6 mm. The substrate extends ∼40 mm to the connector, linking via headstage to an Intan acquisition system. A reference electrode was placed in the frontal region of the craniotomy, ∼25 mm from the array. Signals were sampled at 2 kHz.

**Figure 2.**
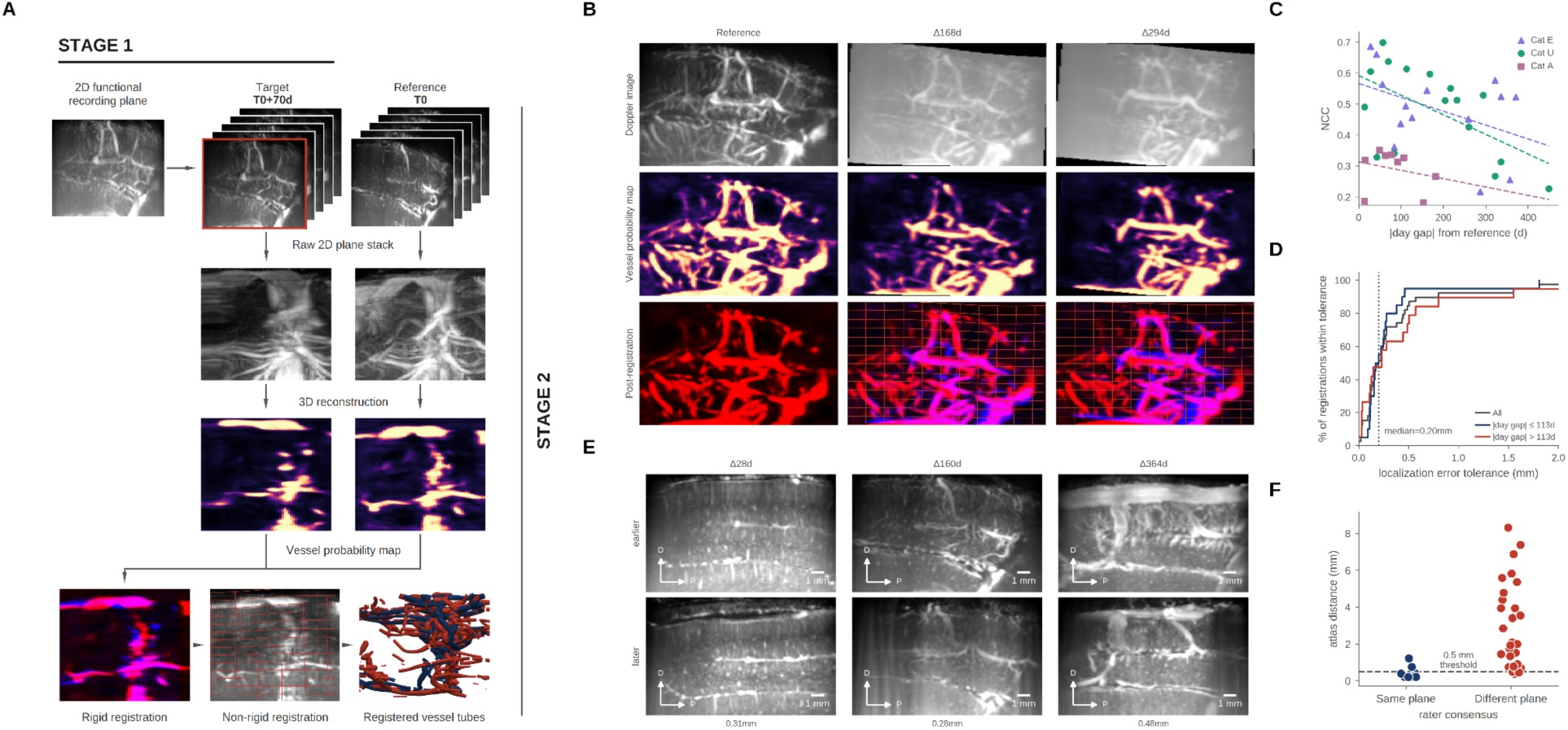
Chronic 3D vessel-map registration provides stable, sub-millimeter anatomical correspondence across day gaps up to ∼450 days. (A) Two-stage registration pipeline. Stage 1: Co-registration of a 2D functional plane to a same-day 3D anatomical volume, followed by transformation to a fixed reference 3D volume to establish a shared atlas coordinate frame. Stage 2: Direct 3D-to-3D registration between reference (T₀) and target (T₀ + 70d) volumes, showing: raw swept 2D sagittal planes, 3D volume reconstruction, vesselFM segmentation, rigid alignment (red = reference, blue = target), non-rigid B-spline warp, and final 3D vessel-tube rendering. (B) Longitudinal registration at the sagittal sinus/midline plane across increasing time gaps (Δ168d, Δ294d). Top: Raw Doppler images. Middle: vesselFM probability maps. Bottom: Post-registration overlays (magenta indicates voxel overlap) with B-spline displacement grids. (C) Post-registration structural similarity (normalized cross-correlation, NCC), computed over the whole 3D volume, versus elapsed time (n = 39 registrations across 3 animals). Dashed lines denote per-animal linear fits from a linear mixed-effects model. (D) Cumulative localization accuracy, assessed at the sagittal midline plane, across early (≤113d, n = 20) and late (>113d, n = 19) registration intervals (n = 39 total). Median localization error is 0.20 mm. (E) Vascular alignment across representative same-tissue pairs spanning day gaps from 28 to 364 days (atlas frame offset: 0.18–0.48 mm). D/P = dorsal/posterior. Scale bar = 1 mm. (F) Blinded manual validation: physical distance to the consensus-matched location for same-pair vs. different-pair candidate matches (n = 41 candidate pairs total), rated blind to session identity by trained raters, with anchor and catch trials included to monitor rating quality.

**Figure 3.**
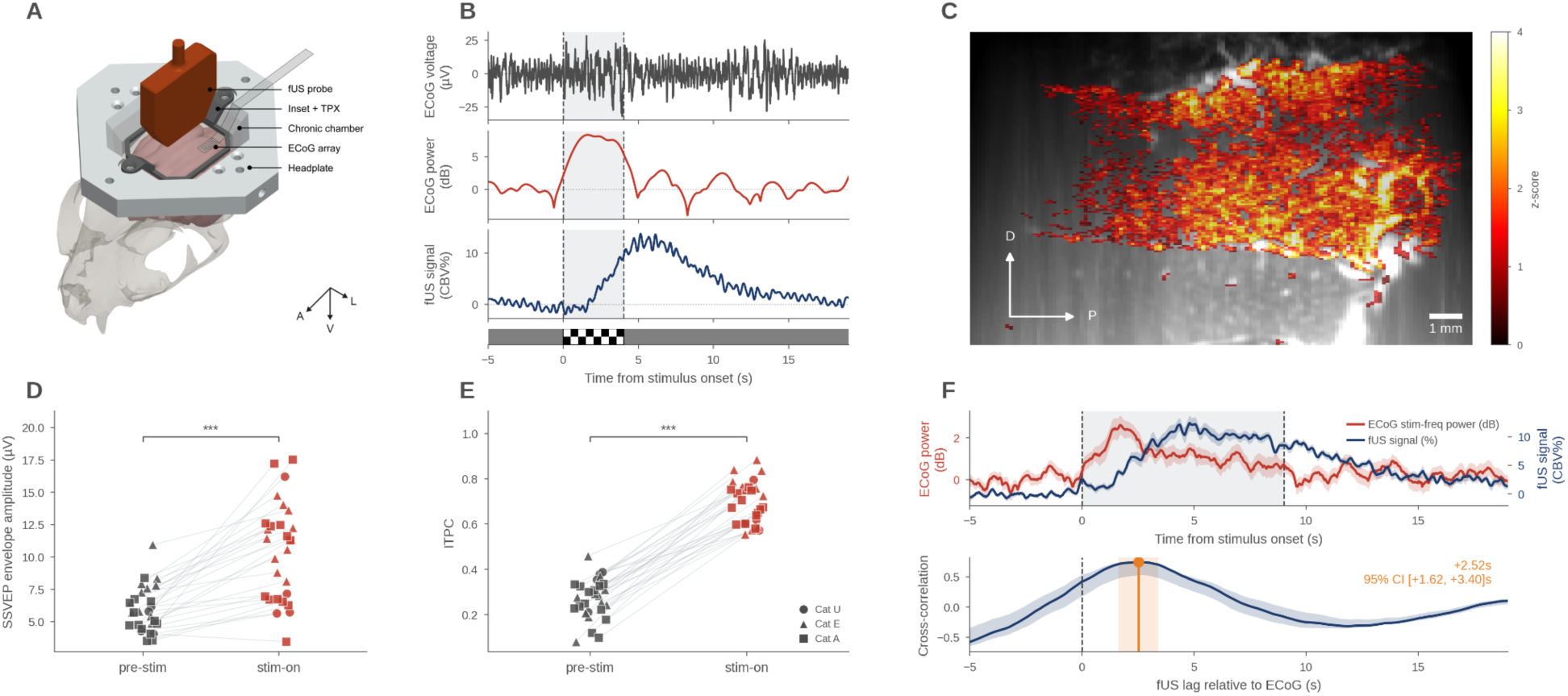
Simultaneous ECoG–fUS recording of visually evoked responses using the chronic multimodal platform. (A) Schematic of the multimodal chamber assembly showing acute ECoG placement under agarose, a TPX-windowed inset mounted to the headplate, and a motorized holder for fUS probe coupling. (B) Trial-averaged (n = 10) multimodal responses to a 4-s, 7-Hz flickering checkerboard (shaded region) stimulation. From top to bottom: broadband ECoG (1–40 Hz), 7-Hz narrowband power change (dB), fUS active-pixel CBV change (ΔCBV%), and stimulus protocol. (C) Spatial fUS activation map (z-score, FDR-corrected q < 0.05) overlaid on anatomical background. (D) SSVEP envelope amplitude and (E) inter-trial phase coherence (ITPC) at baseline vs. stimulus-on (n = 28 recordings across 3 cats; shapes denote animal identity) (F) Neurovascular coupling summary (n = 28). Top: Group-averaged mean ECoG power change (red) and fUS ΔCBV% (blue) time courses (±SEM). Bottom: Cross-correlation between ECoG and fUS signals, showing a well-defined hemodynamic lag peak, with fUS lagging ECoG. D, dorsal; P, posterior; A, anterior; V, ventral; L, lateral. (Significance level: *** p < .001.)

#### Synchronization

A TTL pulse from the stimulus PC, routed through a LabJack to an analog input channel of the Intan acquisition system (sampled at 2 kHz), marks the onset and offset of each visual-stimulus block on both the ECoG and fUS timelines, aligning both signals to within ∼50 ms for trial-locked and cross-correlation analyses (Fig. 3B,F).

#### Visual stimulation

Stimuli were generated in custom Python built on PsychoPy (Peirce, 2007) and presented on a 55-inch OLED display ∼43 cm from the eyes. For activation mapping: full-screen contrast-reversing checkerboard, 0.125 cycles/degree, nominal 7 Hz alternation (true display-refresh-limited ∼6.7 Hz; all frequency-domain analyses used each recording’s own measured frequency), 4 s on, flanked by gray baselines (5 s pre-, 10 s post), 10 repetitions averaged per recording (Fig. 3B). For retinotopy: an 8°-wide contrast-reversing checkerboard bar (0.125 cycles/degree, 6 Hz reversal) swept continuously without intervening baselines; elevation mapping swept bottom-to-top at 3.8°/s, 5 repetitions. Widefield and fUS retinotopy used identical stimulation, so the maps compared in Fig. 4D–F differ only in recorded signal and analysis.

**Figure 4.**
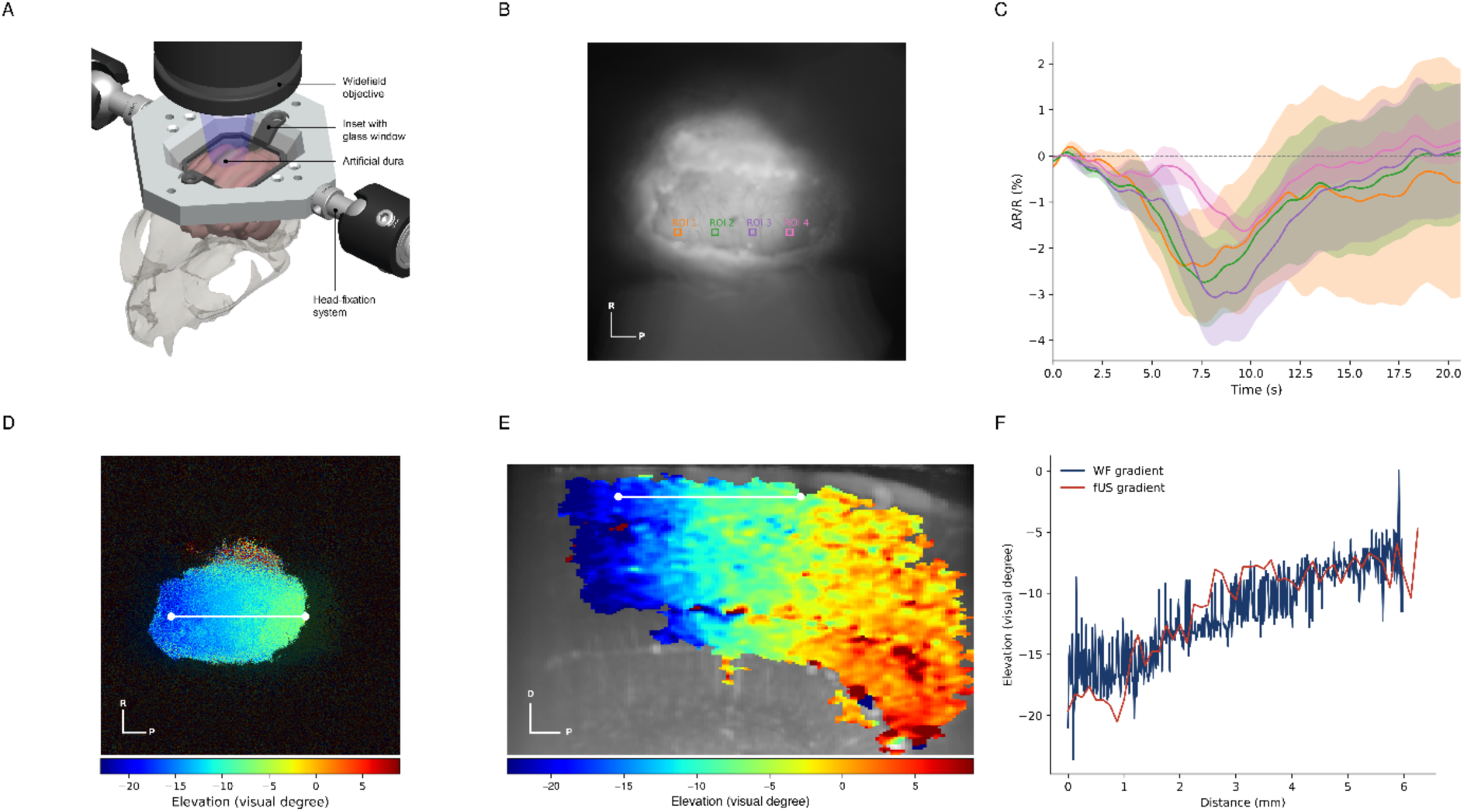
Multimodal retinotopic mapping links intrinsic signal optical imaging and functional ultrasound in cat visual cortex (A) Schematic of the optical imaging chamber assembly. (B) Maximum intensity projection image of a functional mapping intrinsic signal optical imaging recording, with regions of interest (ROI 1-4) indicated. (C) Mean signal traces of the ROIs marked in (B), expressed as percentage change in reflectance (ΔR/R). Shading around curves shows standard error of the mean (SEM). (D) Elevation retinotopic map of cat visual cortex recorded using widefield intrinsic signal optical imaging (WF), reconstructed from the recording shown in (B). Color bar indicates corresponding elevation position in the visual field. (E) Functional ultrasound imaging (fUS) elevation retinotopic map of cat visual cortex, recorded in the same cortical region as (D), overlaid on structural power Doppler image. (F) Comparison of retinotopic gradients extracted along the white marker lines in (D) (WF) and (E) (fUS). The two gradients were significantly correlated in this single representative case (Pearson r = 0.837, permutation test with circular shifts to account for spatial autocorrelation, p < 0.001, 5,000 permutations), confirming close spatial alignment between the two imaging modalities. The direction indicators in (B), (D) and (E) also serve as scale bars; each arm represents 1 mm. D, dorsal; P, posterior; R, right.

### Cross-day anatomical registration

#### 3D scan registration

Individual 3D anatomical scans, each assembled from one session’s swept 2D planes, are registered into an animal-specific reference frame; one session per animal serves as reference, and all others are registered to it independently (Fig. 2A,B). Rigid registration: volumes are converted to vessel-probability maps with vesselFM and aligned by a 3D Euler transform; registration uses this representation rather than raw intensity, whose session-to-session Doppler-contrast variation produces poor local minima. Non-rigid registration: starting from the rigid solution, a cubic B-spline free-form deformation is fit to raw intensity, absorbing residual local misalignment such as minor tissue shift and subtle vascular remodeling.

#### Quality control and validation

Registration quality is tracked over time using two independent measures, each targeting a different failure mode. First, structural similarity — normalized cross-correlation (NCC) between the fully registered (rigid + B-spline) target and reference vessel-probability volumes, computed over the whole 3D volume — is modeled against elapsed time using a linear mixed-effects model with a random intercept per animal (REML; Fig. 2C). Second, localization accuracy is assessed specifically at each registration reference midline (sagittal sinus) plane. An independent ground truth for that plane’s position along the sweep axis in the target volume is obtained by 3D phase cross-correlation between the two vessel-probability volumes, each cropped symmetrically to a common slice count centered on the sweep. This ground-truth position is compared against the position the composed rigid + B-spline transform predicts for the same plane’s center point, giving a physical-distance error; because phase cross-correlation is algorithmically distinct from the mutual-information-based registration, the check does not validate the method against itself. Registrations were split at the sample’s median day gap (113 days) and the two resulting error distributions compared by Mann–Whitney U (Fig. 2D).

#### Placing recordings in the atlas frame

Each functional recording is a single 2D plane acquired separately from its same-day 3D sweep, so its matching plane is found by slice search: all planes are screened by translation-only phase cross-correlation; the five best candidates are refined by rigid registration (translation, rotation, uniform scale; normalized cross-correlation); the highest final correlation is retained. Composing this transform with the session-to-atlas transform (rigid component inverted analytically, B-spline numerically by fixed-point iteration) gives an atlas coordinate, summarized by the recording center point.

#### Cross-day matching

Correspondence between recordings from different days is established by nearest-neighbor search in the shared frame, per animal, excluding same-day neighbors as trivially near-identical. Candidate matches are defined as the nearest different-day neighbor within 0.5 mm and are subsequently tested against human ratings (below).

#### Blinded identity validation

A stratified sample of candidate cross-day pairs — balanced across atlas-distance bins and animal, with anchor and catch trials of known ground truth interspersed — was rated by three raters blinded to session identity, each viewing two anonymized structural images per pair in randomized order with no distance, similarity, or identity information. Ratings used a 4-point ordinal scale (definitely/probably same or different tissue) plus “cannot assess”; only “definitely” ratings were retained, and pair consensus taken as the median of contributing ratings. Agreement was quantified by Krippendorff’s α and pairwise quadratic-weighted Cohen’s κ. The pipeline’s call was evaluated against consensus as sensitivity and specificity (consensus “same tissue” as positive class), and distance to the consensus-matched location compared between same- and different-tissue candidates by one-sided Mann–Whitney U (Fig. 2F).

### Analysis

#### fUS activation mapping

Activation maps were generated by computing per-pixel Pearson correlation between the mean-of-repetitions time course and a canonical hemodynamic-response-function model of the stimulus; one-tailed t-test on the correlation coefficient (positive activation), Benjamini–Hochberg FDR correction (q = 0.05). Responses are expressed as fractional cerebral blood volume change (ΔCBV%) against a pre-stimulus baseline and maps as z-scores against the pre-stimulus noise distribution, displayed on a fixed color scale over each recording’s own anatomical Doppler image (Fig. 1D, 3C, 5A; Dizeux et al., 2019).

**Figure 5.**
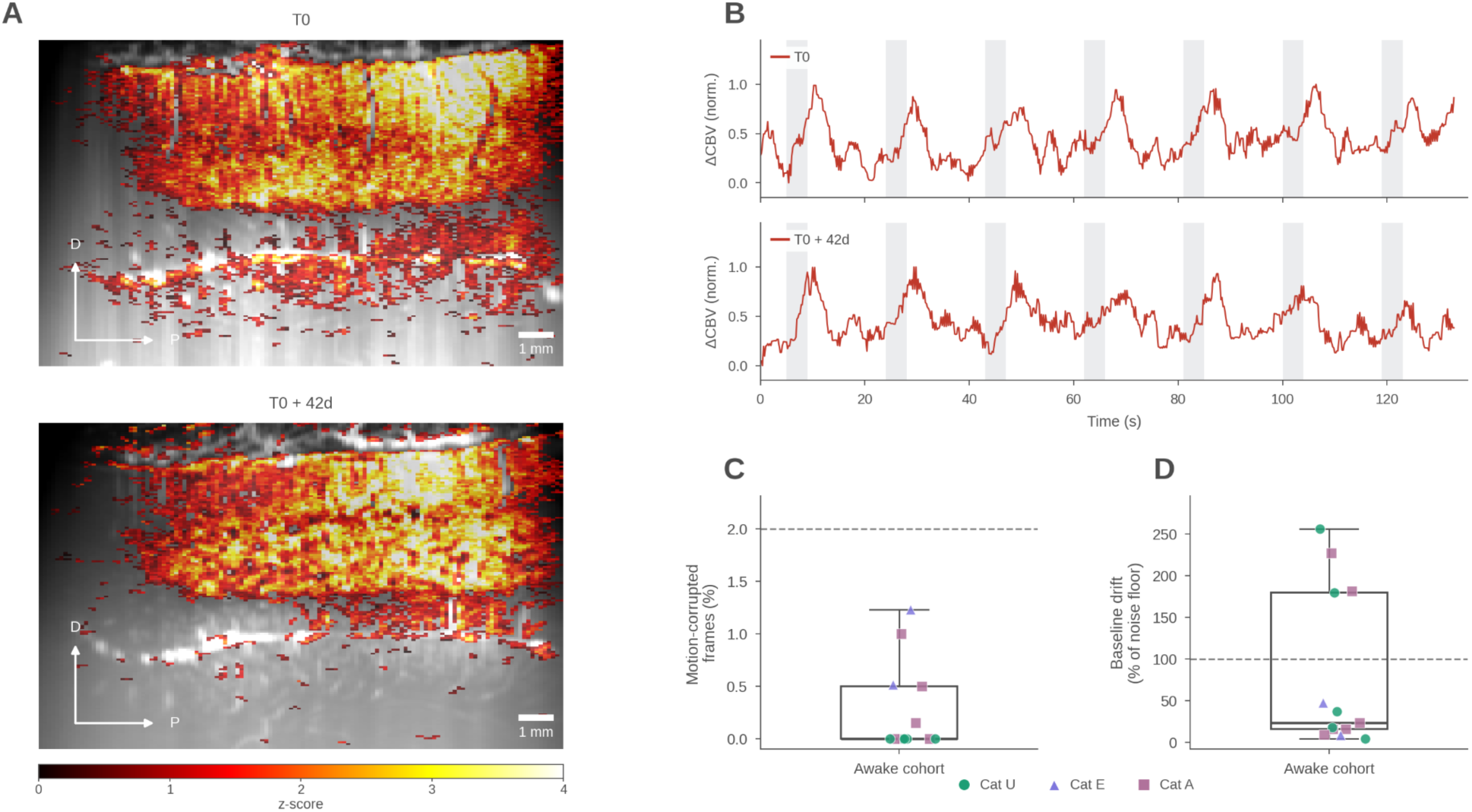
Awake fUS imaging (A) Representative fUS activation maps for two awake recordings from the same cat, 42 days apart, acquired from the same cortical region. D, dorsal; P, posterior. (B) Full-session normalized ΔCBV time course for the two recordings presented in (A) (grey bands mark stimulus period) (C) Motion-corrupted frame fraction in awake cohort (n = 13 recordings, 3 animals) (D) Baseline drift for the same cohort, relative to each recording’s own trial-to-trial response variability, against a 100%-of-noise-floor threshold; In (C, D), marker shape/color indicates animal identity (circle, Cat U; triangle, Cat E; square, Cat A); boxes show median and interquartile range. D, dorsal; P, posterior.

#### fUS retinotopy

Retinotopic maps were derived from continuous periodic visual stimulation, analyzed in the Fourier domain (Kalatsky & Stryker, 2003). Frames were first spatially smoothed with a 3×3 boxcar filter. A per-pixel Fourier transform was then computed over the full recording to extract the power and phase at the stimulus repetition frequency, and phase was converted to visual degrees using the screen dimensions and viewing distance. A pixel was included in the map if its stimulus-frequency power exceeded 1% of the DC power; the map was then restricted to its largest contiguous suprathreshold region, and any holes within that region were filled.

#### Awake data-quality metrics

Two data-quality metrics were evaluated against thresholds set a priori: the fraction of motion-corrupted frames (threshold, 2%) and baseline drift, defined as the deviation of the pre-and post-stimulus baseline from the recording’s own mean, expressed relative to the block-to-block variability of the evoked response (threshold, 100% of this noise floor). Each recording’s metric was tested against its threshold with a one-sided one-sample Wilcoxon signed-rank test, and exact binomial confidence intervals are reported for the fraction of recordings meeting each threshold (Fig. 5C, D).

#### ECoG processing

Channels were cleaned in sequence: robust (MAD-based) thresholding for large-amplitude artifacts, with linear interpolation of flagged samples; a Hampel filter for residual transients; a zero-phase Butterworth high-pass for drift; mains notch filtering; and, common-average referencing across contributing channels. Entrainment was then measured in a narrow band centered on each recording’s own measured stimulus frequency, using a zero-phase bandpass filter followed by a Hilbert transform to extract instantaneous phase and envelope amplitude: SSVEP envelope amplitude and inter-trial phase coherence (ITPC, the magnitude of the trial-averaged unit phase vector) at the stimulation frequency were compared between pre-stimulus and stimulus-on windows (Fig. 3D,E).

#### Widefield retinotopy and segmentation

Frames were averaged across sweeps, and each pixel’s mean response time series low-pass filtered (second-order Butterworth, 3.12 Hz cutoff, zero-phase forward– backward). Retinotopic position is the time of the response extremum, converted to visual degrees from stimulus velocity, screen dimensions, and viewing distance (Fig. 4D). ROI signals are percent change (ΔR/R) against the mean of each sweep’s first 10 frames, with SEM across repetitions for display (Fig. 4C). Two segmentation steps define the analyzed region: tissue is separated from background by mean-intensity projection, rolling-ball background subtraction, light Gaussian smoothing, and Triangle thresholding, with morphological closing and hole-filling to retain one contiguous mask; within it, an activity mask combines the temporal standard deviation of the recorded signal with an 8-neighbor local spatial correlation across time (following CaImAn; Giovannucci et al., 2019), each percentile-normalized, multiplied, re-normalized, and thresholded (Fig. 4D).

#### Cross-modal gradient comparison

One-dimensional elevation profiles were extracted along a common line in each modality (Fig. 4D,E). The medio-lateral position of this line was fixed anatomically: at the start of each fUS session an anatomical scan spanned the full accessible medio-lateral extent, the midline plane was identified from large superficial vessels, and the offset between functional plane and midline computed; in the widefield session, whole-window images located the same midline and the field of view relative to it, and the corresponding line was selected using the fUS-derived offset. Antero-posterior extent was set independently per modality from the elevation maps themselves. Elevation was sampled along the line by linear interpolation against physical distance, and the two profiles resampled onto a common 500-point grid spanning their overlap. Similarity is Pearson’s r; because profile samples are spatially autocorrelated, significance was assessed by a circular-shift permutation test preserving each profile’s autocorrelation (one profile held fixed, the other circularly shifted by a random non-zero offset, 5,000 iterations), with a two-sided p-value as the proportion of null correlations at least as large in absolute value, α = 0.05 (Fig. 4F).

### Statistics

Session- and animal-level comparisons use linear mixed-effects models with cat identity as a random effect (Measurement ∼ Condition + (1|CatID)) to avoid pseudoreplication across repeated sessions within an animal: pre- versus stimulus-on SSVEP amplitude and ITPC (Fig. 3D,E), and post-registration NCC against elapsed time (Fig. 2C). Awake-cohort data-quality metrics (motion-corrupted frames, baseline drift; Fig. 5C, D) were each tested against an a priori threshold using a one-sided one-sample Wilcoxon signed-rank test, with exact binomial confidence intervals reported for the fraction of recordings meeting each threshold. Mann–Whitney U tests were used for the two right-skewed distributional comparisons: early- versus late-interval localization error (two-sided; Fig. 2D) and same- versus different-tissue distance to the consensus-matched location (one-sided; Fig. 2F).

### Code and data availability

The analysis pipeline, CAD files, acquisition and stimulation code, and supporting data will be made publicly available in a versioned repository upon publication, together with documentation sufficient to facilitate reproducibility.

## Results

### A modular implant system delivers stable access across large brain regions and reliable mapping of stimulus-evoked activity via functional ultrasound imaging

To achieve chronic, multimodal imaging, we engineered a modular implant system composed of three structural elements (Fig. 1A): a customized, 3D-printed chronic chamber, a load-bearing headplate, and interchangeable modality-specific insets. For individualized chamber design, high-resolution 3T MRI scans were acquired pre-operatively to segment individual skull morphology and tailor the chamber geometry (Fig. S1A,B). Co-registration of the structural MRI with a 3D cat cortical atlas enabled precise surgical targeting, confirming optical and acoustic access to visual areas (A17, A18, and A19) within the planned craniotomy window (Fig. 1B, Fig. S1C). Chamber bodies were fabricated via stereolithography, with lower skirts contoured to match individual skull curvature. Biocompatibility and mechanical stability were established by coating the skull-facing surface with dental-adhesive and bone-cement layer, reinforced by dental screws along the anteroposterior axis to facilitate long-term osseointegration. This custom design and the developed surgical pipeline exhibited sustained chronic mechanical stability and tissue tolerance in vivo: implant longevity exceeded 1.5 years in two subjects and reached over 3.3 years in a third, with all animals maintaining functional chambers without requiring surgical intervention or replacement (Fig. S1D). The chamber’s standardized upper geometry anchors a rigid metal headplate designed for head fixation and modular hardware extensions (Fig. 1A; Fig. S2A). This interface accommodates interchangeable insets—such as an ultrasound-transparent polymethylpentene (PMP) foil inset or an optical glass inset for widefield imaging—enabling cross-modal experiments within the same craniotomy window without repeating surgeries. To extend standard planar fUS to 3D volumetric acquisitions, we designed a lightweight (<5% total body weight) motorized scanning module docking onto the metal interface (Fig. 1A). The module houses the transducer probe driven along orthogonal axes by dual miniature linear stepper actuators under open-source microcontroller control. In vivo 3D power-Doppler sweeps conducted along sagittal and coronal axes at a 200 um step size yielded detailed vascular volumes measuring 16x10x12 mm and 16x10x13 mm, respectively (Fig. 1C). Finally, visual stimulation elicited robust, spatially localized hemodynamic responses within the visual cortex, as demonstrated by pixel-wise z-score activation maps overlaid on baseline structural power-Doppler images (Fig. 1D).

### Registration reproducibility: the same cortical volume can be revisited across imaging days

To establish fUS as an anchor modality for chronic longitudinal tracking, we developed a two-stage 3D registration pipeline to align functional planes across extended time intervals (Fig. 2A). In the first stage, single 2D functional planes are co-registered to a same-day 3D anatomical volume, which is subsequently mapped to a fixed 3D reference scan to transform functional data into a unified atlas coordinate space. In the second stage, longitudinal 3D volumes (T₀ + Δt) are registered directly to the reference volume (T₀) using a vessel-probability segmentation model (vesselFM) that guides sequential rigid alignment and non-rigid B-spline warping (Fig. 2A, B). Using this pipeline, anatomical correspondence was maintained across time gaps of up to ∼450 days (Fig. 2B).

To determine whether registration quality degrades over such timescales, we evaluated structural similarity (normalized cross-correlation, NCC) as a function of elapsed time using a linear mixed-effects model. This revealed a small but statistically significant decline (≈−0.05 NCC per 100 days; p = .0016; Fig. 2C), consistent with subtle long-term drift in baseline tissue contrast.

Critically, this drift in raw structural similarity did not translate into any loss of spatial precision. Plane-localization accuracy remained stable across the entire recording timeline (Fig. 2D): across the dataset (n = 39 registrations, 3 animals), the pipeline achieved a median localization error of 0.20 mm, with 72% of registrations aligning within a single scan step (0.2 mm) and 97% falling within 1.0 mm. Directly comparing early (≤113 days) against late (>113 days) intervals confirmed no degradation in accuracy over time (Mann– Whitney U test, p = .870; Fig. 2D). This robust spatial correspondence, sustained despite gradual contrast drift, enabled reliable identification of matching microvascular structures across time gaps ranging from 14 to 364 days (Fig. 2E) — establishing that anatomical registration precision is fully preserved even as raw image contrast evolves over months. To independently validate the automated registration pipeline, a blinded manual-rating procedure assessed whether candidate matched pairs identified across days were genuinely the same tissue (Fig. 2F). Untrained raters, blind to session identity and including anchor and catch trials to monitor rating quality, judged candidate pairs against a consensus standard (n = 41 candidate pairs). The automated pipeline’s matches showed high sensitivity (0.80) and specificity (0.84) relative to this consensus. Physical distance to the consensus-matched location was markedly smaller for same-pair than for different-pair candidates (median 0.23 mm vs. 1.93 mm; Mann–Whitney U = 289, p = 2.5×10⁻⁵, rank-biserial correlation = 0.93), providing independent, human-rated confirmation that the pipeline’s automatically identified matches correspond to genuinely same-tissue locations.

### Simultaneous ECoG–fUS recording of visually evoked responses using the chronic multimodal platform

Simultaneous ECoG–fUS recordings were performed during full-screen flickering checkerboard stimulation (7 Hz, 4 s) with the ECoG array placed over the marginal gyrus (gyrus marginalis, area 17). Visual stimulation reliably evoked robust steady-state visually evoked potentials (SSVEPs) across all animals (n = 28 recordings, 3 cats). Relative to pre-stimulus baseline, stimulation significantly increased SSVEP envelope amplitude by 4.51 µV (95% CI [3.06, 5.95], LME model, p < .001; Fig. 3D), and inter-trial phase coherence rose by 0.424 over the same window (95% CI [0.377, 0.471], LME model, p < .001; Fig. 3E), confirming reliable phase-locking to the stimulus across recordings and animals. Cross-correlating the group-averaged ECoG power and fUS CBV% time courses revealed a peak at a fUS lag of +2.52 s relative to ECoG (95% CI [1.62, 3.40] s; Fig. 3F) — consistent with the expected delay of the hemodynamic response relative to the underlying electrophysiological event, and demonstrating that the platform recovers a quantitative, reproducible neurovascular-coupling estimate from concurrent recording through a single chronic chamber.

### Widefield optical imaging recovers a retinotopic map that matches the fUS-derived map

To validate the chamber as an optical imaging port and to cross-validate fUS-based functional mapping against an established modality, we imaged cat visual cortex through the chamber’s glass-window insert using widefield intrinsic signal optical imaging (Fig. 4A,B) and compared the resulting retinotopic organization with fUS maps recorded from the same cortical region. The resulting elevation map spans approximately −20° to −5° of visual elevation across the imaged region (Fig. 4D, shaded by an activity-based segmentation mask). Functional ultrasound imaging of the same cortical region, using the identical stimulation paradigm, reproduced this organization (Fig. 4E). Despite being derived from cerebral blood volume changes rather than activity-dependent surface reflectance, the fUS elevation map showed the same smooth, near-monotonic progression of preferred elevation along the antero-posterior axis, over a comparable cortical extent and range of visual degrees (approximately −20° to −5°).

To quantify this correspondence, we extracted one-dimensional elevation profiles along lines placed at the same medio-lateral cortical position in both modalities, anchored anatomically to the cortical midline identified from large superficial vessels (see Methods), and resampled them onto a common 500-point grid. The widefield and fUS gradients were strongly correlated (Pearson r = 0.837; Fig. 4F). Because neighboring samples along each profile are spatially autocorrelated, significance was assessed with a circular-shift permutation test that preserves each profile’s autocorrelation structure; the observed correlation clearly exceeded the resulting null distribution (p < 0.001, 5,000 permutations). Two distinct signals therefore report the same retinotopic organization on the same chronically accessible tissue, cross-validating the chamber’s optical and ultrasound modalities within a single preparation.

### Awake fUS imaging is reproducible across weeks

Studying brain function in awake, behaving animals is essential for linking cortical activity to natural behavior and arousal state. We therefore tested whether the chronic chamber and its motorized fUS module could support stable functional imaging in awake cats without any modification to the implant or imaging hardware. Animals were gently body-restrained but left head-unrestrained, since the transducer mounts rigidly to the chamber rather than requiring external fixation.

As a demonstration of this capability, we compared two awake recordings from the same animal and cortical region, acquired under matched visual stimulation on different days. Activation maps recovered highly overlapping suprathreshold regions across sessions (Dice coefficient = 0.76 after co-registration onto a shared pixel frame; Fig. 5A), and the full-session ΔCBV time course showed matching response dynamics across sessions (Fig. 5B).

To characterize data quality across the broader awake cohort, we quantified motion-corrupted frames and baseline drift across multiple awake recordings (n = 13 recordings, 3 animals) against thresholds set a priori. Motion contamination was minimal throughout: all 13 recordings fell below a 2% motion-corrupted-frame threshold (95% CI [75%, 100%]) one-sided Wilcoxon signed-rank test against threshold, p < .001; Fig. 5C), confirming that the skull-anchored transducer maintains imaging-plane stability without head fixation. Baseline drift, expressed relative to each recording’s own trial-to-trial response variability, was more variable across the cohort: 9 of 13 recordings (69%, 95% CI [39%, 91%]) fell within a threshold of 100% of this noise floor, and as a group the cohort did not differ significantly from the threshold (p = .227; Fig. 5D). The wide confidence interval reflects a small number of recordings with drift substantially exceeding the response’s own noise floor, rather than a uniformly elevated baseline across the cohort; such drift is consistent with the arousal- and behavior-linked hemodynamic fluctuations documented in awake recordings generally, discussed further below.

## Discussion

We have described a modular chronic cranial platform that provides individualized, longitudinal access to a large cortical region in the cat and accommodates functional ultrasound, widefield optical imaging, and electrocorticography through a single craniotomy. Four claims follow from the data. First, a customized, 3D-printed chamber can remain mechanically and functionally stable for years, a durability previously achieved only with machined metal implants. Second, the ability to re-identify a specific cortical volume across months or even years, was established for fUS, the anchor modality throughout. Third, the modular interface converts the same craniotomy between hemodynamic, optical, and electrical readouts without further surgery, including concurrent ECoG and fUS from the same tissue volume. Fourth, the platform supports functional imaging in awake, head-unrestrained animals, extending longitudinal access beyond rigid head-fixed preparations.

### Current bottlenecks in chronic access

Chronic imaging traditionally confronts a trade-off between structural durability and geometric flexibility. Machined titanium or stainless steel chambers achieve multi-year mechanical stability but are difficult to customize to complex, individual skull topologies. The present platform resolves this specific trade-off: customized, low-profile 3D-printed resin chambers combine individualized geometric fitting with multi-year mechanical durability (>3.3 years).

A separate, longstanding limitation of chronic optical windows is degradation of the interface itself — dural fibrotic thickening, tissue regrowth, or membrane clouding — which progressively restricts optical or acoustic access even when the surrounding chamber remains structurally sound. The present design addresses this through an artificial dura intended to limit dural regrowth at the tissue interface, though its long-term optical performance over multi-year timescales was not separately quantified in this study.

Another pervasive challenge in longitudinal preparations is distinguishing true physiological or plastic remodeling from spatial misalignment across sessions. Conventional longitudinal studies frequently rely on “implant survival” as a surrogate for signal stability; however, subtle tissue shifts and dural re-vascularization introduce unquantified spatial variance that blurs longitudinal metrics. By implementing an automated vascular registration pipeline, we demonstrated sub-millimeter localization error (0.20 mm median) across intervals reaching ∼450 days. Registration precision did not decay over time, even as whole-volume structural similarity declined modestly, indicating that local vascular geometry preserves landmark coordinates independently of global signal shifts. In practice, this converts each subject into its own anatomical reference across months or years, removing a major source of between-session variance and enabling experimental designs that track specific circuits over time.

### Multimodal capabilities and current limitations

Architecturally, the chamber converts the same tissue volume between hemodynamic, optical, and electrical readouts without secondary surgery. In single-session cross-validations, intrinsic signal optical imaging and fUS produced concordant retinotopic maps (r = 0.837, p < .001) despite relying on distinct contrast mechanisms—activity-dependent surface reflectance versus cerebral blood volume. However, unlike fUS, the longitudinal stability of widefield optical imaging was not evaluated in the present study. Extending optical imaging to comparable multi-month timescales therefore remains an important objective and will require refined dural management to preserve optical clarity at the cortical surface.

The same multimodal architecture also permits electrophysiological cross-validation. An ultra-thin, ultrasound-transparent polyimide ECoG array was positioned directly on the marginal gyrus while fUS simultaneously scanned the underlying cortical volume through the array. The polyimide array produced no detectable acoustic shadowing or imaging artifacts in the fUS images. The combined configuration yielded consistent stimulus-evoked responses across modalities, with repeatable, stimulus-specific electrophysiological signatures in the ECoG and a highly reproducible hemodynamic lag in the corresponding fUS response.

ECoG was intentionally deployed in acute paradigms here to establish cross-modal correspondence rather than to assess chronic electrophysiological stability. Because the polyimide array preserves acoustic transmission without degrading fUS image quality, this interface could be incorporated into the chamber for chronic implantation (Panskus et al., 2026). Chronic integration of ECoG with longitudinal fUS would extend the platform toward direct, long-term characterization of neurovascular coupling across graded anesthesia depths—a factor that has long confounded hemodynamic readouts (Masamoto & Kanno, 2012). In such experiments, electrophysiological state markers such as 1/f spectral slope and Lempel–Ziv complexity could provide independent measures of cortical excitation–inhibition balance (Medel et al., 2023), allowing changes in hemodynamic responses to be interpreted against the underlying neural state.

### Awake acquisition and physiological baseline variability

Because the ultrasound transducer docks directly to the skull-anchored chamber, its position relative to the underlying cortex is maintained without requiring rigid head fixation. Motion contamination in awake animals was minimal (<2% corrupted frames), and functional topography remained highly stable across a 42-day interval (Dice = 0.76). Nevertheless, approximately one-third of awake sessions exhibited baseline hemodynamic drift exceeding the sensory response’s noise floor. Rather than reflecting a chamber-specific flaw or mechanical instability, these slow fluctuations mirror established physiological sources. Awake recordings carry large, state-dependent hemodynamic shifts tied to arousal, facial movement, and vigilance transitions, which frequently exceed evoked response amplitudes across awake imaging modalities generally (Zhang et al., 2022). Similar findings in longitudinal awake fUS demonstrate that functional readouts are inherently less reproducible across sessions than structural or blood-flow metrics, with habituation stress serving as a primary driver (Huang et al., 2026). Explicitly incorporating behavioral covariates—such as pupil dilation, whisking, or locomotion—directly into the fUS general linear model, rather than treating drift purely as noise, provides a proven strategy to recover clean stimulus-evoked signal (Qin et al., 2026). The value of awake acquisition lies precisely in capturing this physiological state space; future paradigms can systematically monitor and model this variance rather than suppressing it through anesthesia.

## Conclusion

Together, these results redefine the operational standard for chronic preparations, shifting the benchmark from passive implant survival to quantified, sub-millimeter spatial re-assessment. Establishing verifiable long-term accessibility to identical cortical volumes opens new avenues for investigating processes such as sensory deprivation, learning, and targeted neural perturbation within individual subjects. Furthermore, the core architectural principle—durable, customizable, and non-disruptive acoustic access—extends well beyond animal research. The recent deployment of permanent cranial windows for repeated fUS monitoring in awake human patients outside the operating room (Rabut et al., 2024) highlights how individualized acoustic interfaces can bridge preclinical innovation and emerging translational paradigms.

## Acknowledgements

The authors gratefully acknowledge all colleagues whose contributions supported the successful completion of this work. We are particularly grateful to Fanni Somogyi and Beatrix Kovács for their extensive support with surgical procedures and experimental work. We sincerely thank Sarolt Kinga Gintner, Renáta Radics, Luca Benedek, Zsombor Fülei, and Hanna Orvos-Nagy for their dedicated assistance with animal care. We also thank Gergely Márton and István Ulbert for their support.

This research was supported by grant 2019-2.1.7-ERA-NET-2021-00047, the Lendület (“Momentum”) Programme of the Hungarian Academy of Sciences, Excellence 151368 funded by the Ministry for Innovation and Technology of Hungary through the NRDI Fund, CELSA/24/020, and KSZF-161/2024 awarded to DaH. The 2024-2.1.2-EKÖP-KDP University Research Scholarship Program – Cooperative Doctoral Program of the Ministry for Culture and Innovation, funded through the National Research, Development and Innovation Fund, was awarded to KCs at Semmelweis University.

## Supplementary Information

### Supplementary Protocol S1 — Full chamber implantation and anesthesia protocol

This protocol provides the complete, step-by-step surgical and anesthesia detail summarized in Methods (“Surgical implantation and chronic maintenance”), for labs seeking to reproduce the procedure directly. Chamber implantation proceeds in three staged surgeries — outer chamber implantation, craniotomy, and durotomy with definitive artificial-dura placement — each separated by a minimum two-week recovery interval. Animals undergo one month of pre-surgical conditioning beforehand (transition to wet food, daily handling to habituate the animal to post-operative care).

### S1.1 Anesthesia and analgesia (identical across all three surgeries)

Animals are premedicated with 0.1 ml/kg maropitant (SC) the afternoon before surgery and fasted ≥ 12 h preoperatively. On the day of surgery: premedication with midazolam (0.25 mg/kg) and medetomidine (0.025 mg/kg), IM; intravenous cannulation followed by chlorphenamine (Suprastin, 1 mg/kg IV); induction with alfaxalone (2–5 mg/kg IV, to effect), ketamine (0.5 mg/kg IV), and fentanyl (0.002 mg/kg IV) — butorphanol is explicitly avoided, as it antagonizes fentanyl. Animals are intubated with a cuffed endotracheal tube (3.0 mm ID adult female, 3.5 mm ID adult male). Neuromuscular blockade (rocuronium, 0.6 mg/kg IV bolus) is induced only after intubation to enable fully controlled ventilation, which contributes to intraoperative edema prevention; furosemide (2 mg/kg IV) is given prophylactically at induction for the same reason. Anesthesia is maintained with continuous-rate infusions of alfaxalone (2–5 mg/kg/h), ketamine (0.6 mg/kg/h), rocuronium (0.48 mg/kg/h), and fentanyl (0.002 mg/kg/h); animals are mechanically ventilated (volume-controlled, 18 breaths/min at baseline, titrated against end-tidal CO₂ if edema develops — see S1.2). At the end of surgery, alfaxalone is stopped at the start of skin suturing to facilitate timely emergence; ketamine, fentanyl, and rocuronium infusions are stopped once the animal is removed from the stereotaxic frame. Neuromuscular blockade is reversed with neostigmine (0.04 mg/kg IV), with atropine (0.02 mg/kg IV) given for bradycardia — always after, never before, neostigmine, to avoid neostigmine-induced bradycardia. Medetomidine sedation is reversed with atipamezole (IM) once ≥ 1 h has elapsed since stopping the alfaxalone infusion. The intravenous cannula is deliberately left in place for at least 24 h post-surgery.

### S1.2 Intraoperative edema management

An anti-edema contingency ladder is applied identically during the craniotomy and durotomy surgeries. If brain swelling is observed intraoperatively: furosemide (4 mg/kg IV) is given first; end-tidal CO₂ is then checked and, if above 35 mmHg, ventilation rate is increased (to 19–22 breaths/min) to bring EtCO₂ down, without allowing it to fall below 30 mmHg; if swelling persists despite these measures, a mannitol infusion (1 g/kg, infused over ≥ 20 min) is started. The same mannitol step is used if swelling is observed after the chamber has been closed. With the staged protocol followed as described, intraoperative brain swelling is not expected to occur.

### S1.3 Chamber fixation (outer chamber implantation)

The chamber position is defined stereotaxically, with the posterior craniotomy edge placed 6 mm behind the interaural line; positioning further posterior is avoided to prevent excessive drilling over the tentorium. After scalp shaving and aseptic skin preparation, the skin and underlying muscle are excised around the chamber footprint under local lidocaine analgesia, with hemostasis by bipolar coagulation. The exposed skull is mechanically cleaned (bone scraper, scalpel) and chemically cleared of residual soft tissue with 10– 30% H₂O₂, then dried and re-cleaned by blade. The bonding surface is prepared with a dental etch-and-prime sequence: 37% phosphoric acid etchant (1 min contact, wiped off), a dentine/enamel primer applied to the skull, a separate two-component primer applied to the chamber’s mating surface, and a UV-curable bonding agent (Adper Single Bond) light-cured across the skull surface. The chamber is then fixed in three sequential steps: (1) a stiff-consistency bone cement (Palacos MV+G) beds the chamber base against the skull before screw placement; (2) four screws per side (2.1 mm pilot holes), spanning the full anteroposterior axis of the chamber, are hand-tightened, explicitly shallow enough to avoid brain contact; (3) a low-viscosity cement seals any residual skull–chamber gap, followed by a thick-consistency cement covering the screws and outer chamber wall. All cement surfaces that will contact skin or muscle are then covered with a hydroxyapatite paste (INNOTERE Paste-CPC) to support soft-tissue integration; where hydroxyapatite is used, the skin is not additionally glued to the chamber (skin-to-chamber cyanoacrylate adhesive is used only when hydroxyapatite is omitted). Skin is closed in two layers (epidermal and intradermal running sutures), and the chamber interior is filled with silicone and sealed with a screwed 3D-printed cap until the craniotomy surgery.

### S1.4 Craniotomy

At least two weeks after chamber implantation, the craniotomy (30 × 30 mm) is performed under the same anesthesia protocol. After re-sterilizing the skin and chamber and removing the interim silicone/cap, the bone within the chamber’s inner perimeter is drilled in three stages of decreasing burr size (> 2.1 mm, then 2.1 mm, then 0.5 mm for the final thin bone layer), beginning along the long edges and proceeding more cautiously over midline areas; a surgical microscope is used for the midline where needed. Continuous sterile saline irrigation and suction are used throughout to limit thermal damage, and local lidocaine is applied continuously. Once the bone segment is mobile, the entire chamber is soaked in sterile 4°C saline for 15 minutes to facilitate dural detachment from the bone before the bone flap is lifted free with two pairs of forceps (requiring two sterile personnel). Bleeding is controlled with bone wax and a haemostatic agent (Lyostypt); bipolar coagulation is used for non-midline bleeding only — coagulation is never used at the midline, since this risks clotting the underlying dural venous sinus and consequent, potentially fatal, brain swelling. For the fUS variant of the chamber, the dura is left intact: if it opens spontaneously during drilling, the opening is covered with sterile Lyostypt; if it remains closed, two small deliberate dural openings are made anterolaterally (bilaterally) and covered the same way. The craniotomy is cleaned and dried, packed with sterile Lyostypt, then filled with silicone and re-sealed with the 3D-printed cap. The edema contingency ladder (S1.2) applies throughout.

### S1.5 Durotomy and artificial dura placement

At least two weeks after the craniotomy, the dura is opened under the same anesthesia protocol. The durotomy margin is kept at least 1 mm from the midline, and the dura is allowed to dry fully before cutting, which facilitates clean separation from the underlying brain surface. The dura is opened with an 18G needle along the planned edge and lifted with fine forceps; where material must be tucked beneath the dura (e.g., the artificial dura sheet itself), blunt fine needle-holders — rather than scissors — are used to free the dural edge from the brain surface without direct trauma to the cortex. The durotomy is closed as pre-planned, covered with sterile Lyostypt, and the chamber is filled with silicone and re-sealed with the 3D-printed cap. The same edema contingency ladder (S1.2) applies.

### S1.6 Post-operative care

On the day of each surgery, the following are administered: tramadol (1 mg/kg SC, analgesia), meloxicam (0.1 mg/kg SC — double the standard maintenance dose, intentionally, as a first-day loading dose), marbofloxacin (8 mg/kg IV, diluted 10× in sterile saline and given over ≥ 5 min), amoxicillin/clavulanic acid (28/7 mg/kg SC), and maropitant (1 mg/kg SC). If spontaneous breathing has not returned within 20 minutes of stopping neuromuscular blockade, end-tidal CO₂ is targeted to 35–38 mmHg; if this does not resolve the issue within a further 5 minutes, the pre-calculated neostigmine dose is given IV, always followed by atropine. Extubation may be delayed by up to 60 minutes relative to the shorter, shallower anesthesia used for recording sessions. Oral gabapentin (4 mg/kg) is given once normal swallowing has returned (typically 2–4 h post-extubation, but potentially longer depending on alfaxalone clearance) and is required before the animal is left unattended. The intravenous cannula is not removed on the day of surgery; removal is deferred to the following day at the earliest, and only once the animal is behaving normally and eating.

---

**Supplementary Figure 1.**
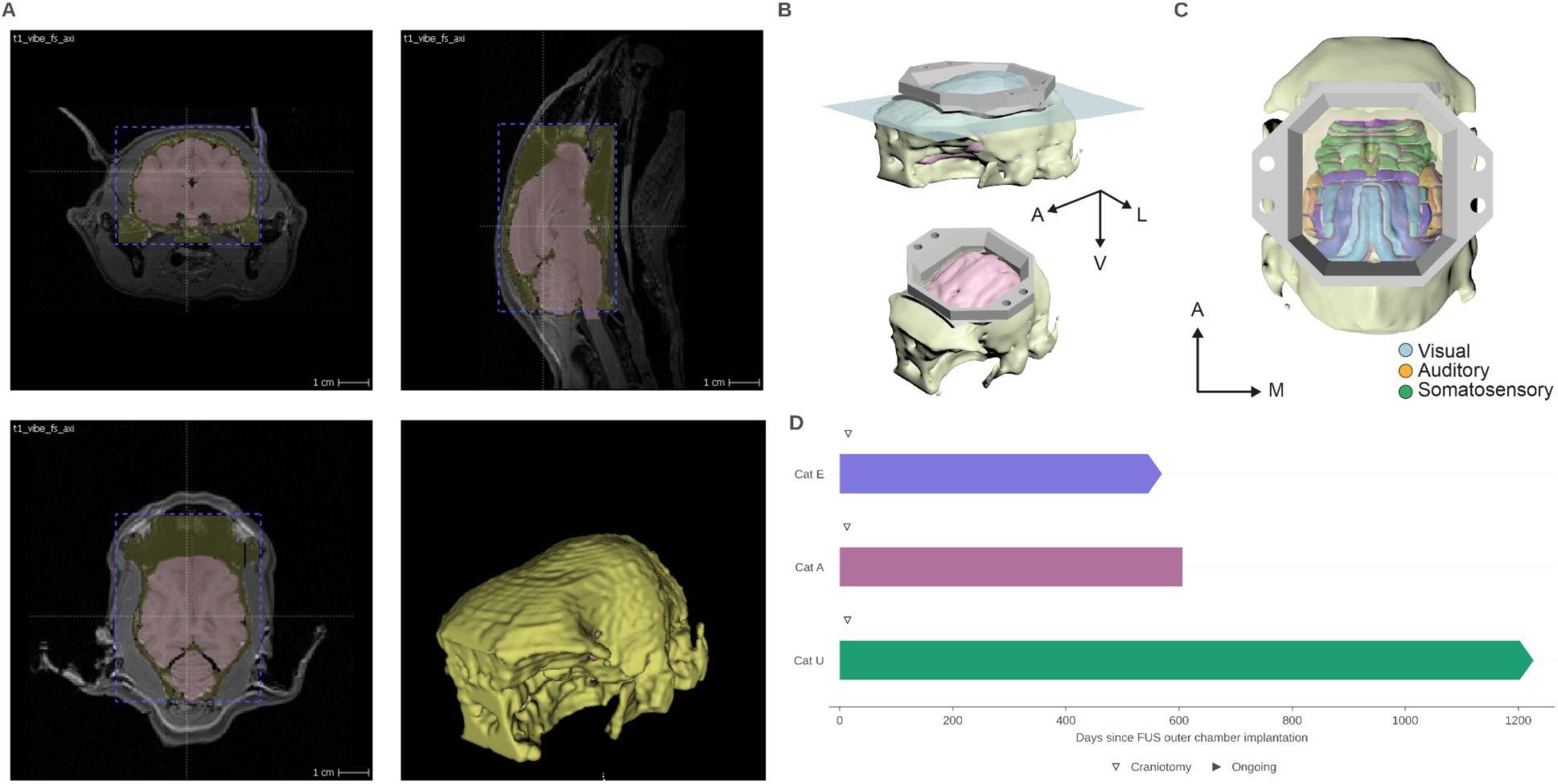
MRI-based individual-animal chamber planning. (A) Automated skull and brain segmentation from a T1-weighted MRI scan. (B) Reference-guided implant fitting and virtual skull cutting used to define the chamber footprint and craniotomy for the individual animal. (C) 3D brain-atlas registration predicting which functional areas (visual, auditory, somatosensory) fall within the planned chamber access before surgery. (D) Chamber longevity for the three implanted animals. Bars indicate the duration of functional chamber access from implantation for the three cats included in this study (Cat E, 570 days; Cat A, 606 days; Cat U, 1227 days); arrowheads indicate chambers that remain in ongoing use.

**Supplementary Figure 2.**
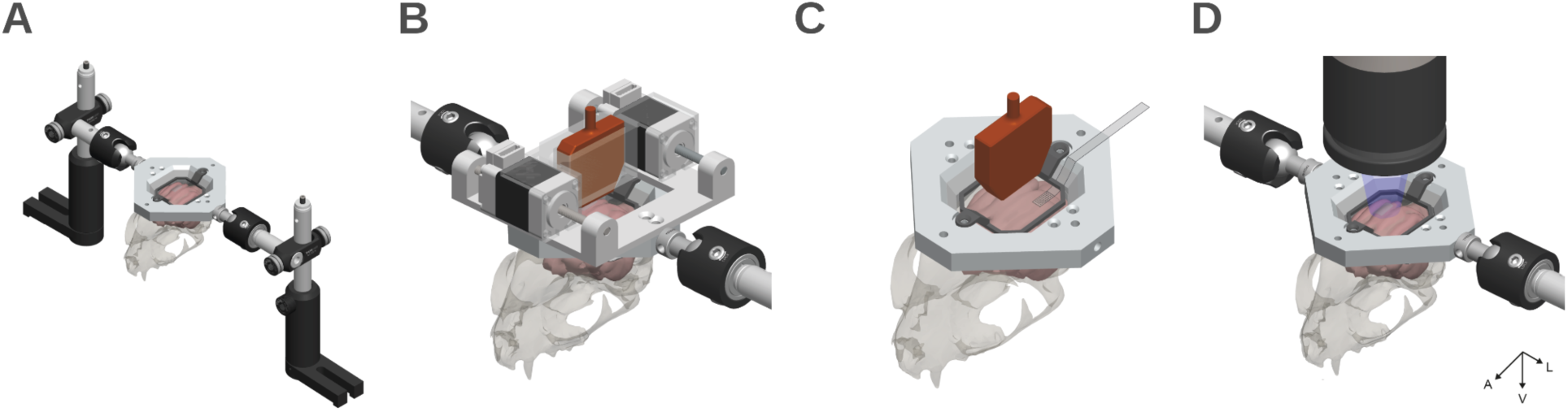
Modular hardware extensions supported by the standardized headplate interface. (A) Head-fixation configuration: the load-bearing metal headplate is clamped bilaterally by post-mounted Thorlabs holders, rigidly securing the head. (B) Motorized fUS scanning module docked to the headplate, housing the ultrasound transducer (orange) driven along two orthogonal axes by miniature linear stepper actuators, with the head-fixation clamps engaged as in (A). (C) ECoG configuration: a thin-film electrode array is introduced through the open chamber onto the cortical surface beneath the transducer position. (D) Widefield optical imaging configuration: objective is positioned above the optical glass inset, with the illumination path directed onto the exposed cortex. A, anterior; L, lateral; V, ventral.

